# Blue light compromises cellular fate of intrinsically photosensitive Müller glial cells in the developing chicken retina

**DOI:** 10.64898/2026.10.26.753828

**Authors:** NA Marchese, MN Rios, A Yaneff, C Davio, ME Guido

## Abstract

Müller glial cells (MCs), the predominant glial population in the retina, interface with all neuronal types to form the structural and functional core of the tissue. Beyond their established homeostatic roles, MCs have emerged as promising endogenous neural stem cells with remarkable regenerative potential. Our recent work identified MCs as novel photosensitive elements of the vertebrate inner retina that express non-visual opsins and respond to brief light stimuli. Direct photic responses manifest as a sustained increase in intracellular calcium levels driven by a G protein–mediated signaling cascade that primarily recruits calcium from the endoplasmic reticulum, highlighting MC photoreception and its roles in retinal physiology. In the present work, we characterize expanded light-directed responses in primary chicken MC cultures, identifying a blue light (BL)–triggered signaling cascade. BL stimulation induces a peak in cAMP levels at 10 min, followed by CREB activation (p-CREB) within 30 min—a temporal profile consistent with observations in *ex vivo* retinal explants. BL exposure (1 h) at the onset of each cell cycle dampens proliferative capacity and migration while lengthening the cell cycle in both primary chicken MCs and an immortalized human MC cell line compared to dark controls. Furthermore, over time (24, 48, and 72 h), BL exposure reduces the proportion of pluripotent cells (Sox2 positive cells), concomitant with an increase in neuronal progenitors (Tbr2) and no change in glial markers (GS, RGR, GFAP). Finally, mature neuronal markers (TUJ-1 and Brn3) increase in MC cultures 120 h post-BL stimulation. Overall, this work broadens our understanding of BL-triggered signaling cascades in MCs and uncovers photoreception as a potential mechanism driving cell fate in the developing retina.

## Introduction

Photoreception evolved as a reliable mechanism to capture ambient light and guide diverse molecular, cellular, and physiological processes ^1^. At the molecular level, light perception relies on a G-protein-coupled receptor (the apoprotein) bound to a light-sensing chromophore derived from vitamin A. These photopigments, which share an evolutionary origin with melatonin receptors, diverged from a common ancestor into four major clades: one restricted to cnidarians and three present in bilaterians, which have expanded into a vast array of opsins—some of which remain undescribed ^2^. Consequently, vertebrate opsins govern both visual and non-visual functions across ocular and extraocular tissues, frequently conserving wavelength sensitivity in the blue/violet spectrum ^3–6^ .

Upon photon incidence, the chromophore undergoes isomerization from a *cis* to an *all-trans* state, resulting in a conformational change in the opsin that activates and, therefore, dissociates the G-protein to interact with downstream second messenger systems ^2,7,8^. Opsins couple to at least three G-protein subclasses: Gɑi/o proteins, specifically transducin (Gɑt) producing a transient decrease in cyclic nucleotide levels; Gɑq protein activating pathways involves phosopholipase C (PLC) activation; and Gɑs protein, which activates adenylate cyclase. Furthermore, some opsins do not interact with any G protein and do not initiate cell signalling at all (for instance, the retinal G protein-coupled receptor, RGR, playing a role in 11-cis-retinal synthesis and recycling) ^9,10^. Overall, as a larger diversity of extra-retinal opsin G-protein binding is investigated, more diversity in signal transduction is being continuously found. Given these facts, a G-protein-based opsin classification scheme still lacks confidence for any underlying co-evolutionary pattern to truly understand how opsin signalling has evolved ^10^.

The vertebrate retina, particularly that of birds, contains several different types of photoreceptor neurons: classical photoreceptors, rods and cones; and non-canonical ipRGCs and horizontal cells expressing Opn4 ^2,9,11,12^. In recent years, we have further contributed to the identification of novel non-neuronal photoreceptors by showing that the most abundant retinal glial cell type, Müller glial cells (MCs), express non-visual opsins (sensitive to the blue-UV light range) and respond to short blue light stimuli. Our pioneering evidence identified the expression of RGR, Opn3, and Opn5 in MCs primary cultures derived from the inner retina of birds ^9,13^. In particular, for Opn3, a stepwise increase in its levels was observed throughout development until birth, coinciding with the rise of classical glial markers. Moreover, Opn3 expression in MCs in cultures is regulated by blue light (BL), increasing its expression and altering its cellular localization after 1 hour of light exposure, both events being part of a single response and dependent on protein synthesis ^13^. Subsequently, we identified a direct photic response by MCs specifically to a BL pulse, observed as a significant increase in intracellular calcium levels sustained over several minutes. Notably, this response was seen in a subpopulation of cultured MG (50% of the cells analyzed) and is dependent on opsin activation ^13^. Through more recent studies, we described the BL-triggered G-protein signaling cascade driving calcium mobilization from the endoplasmic reticulum ^14,15^.

MCs, unlike other retinal glial components, are derived from neural crest cells showing a developmental program highly conserved across species ^16^. MCs undergo substantial morphological changes across the different developmental stages up to the acquisition of the complete functional capabilities in the adult retina, where they constitute unique functional elements that span the whole structure ^16–18^. This particular anatomical position allows them to interact with all neuronal types and, therefore, to play key roles in retinal homeostasis, neurotransmission, chromophore recycling, responses to injury, light detection, and neuronal regeneration ^15,19,20^. This capability has been extensively studied in different neurodegenerative models, indicating that MCs are the only cell type in the mature retina with the potential to re-enter cell cycle, proliferate, de-differentiate and restore neuronal loss ^17^. Nevertheless, and despite several conserved features of MCs across different species, this particular capability is restricted to some vertebrates, being fully functional in teleosts, partially active in avian species and showing very limited potential in mammals ^15^. The challenge remains in identifying the mechanisms that hinder regeneration as well as new key stimuli that could uncage the regenerative potential in these species.

The newly discovered intrinsic photosensitivity of MCs raises new questions about the effects of BL stimulation in MCs considering their possible cellular responses, their functionality and the tissue-level activity of this property along developmental stages. Therefore, the present work aimed to further inquire about the BL-driven responses in MC primary culture considering the signalling cascade and the consequences over MCs features, unmasking a previously unrecognized potential effect of light in driving cellular fate in the developing retina.

## Materials and Methods

### Animal Handling

For the different studies performed, chicken embryos (Gallus gallus domesticus), raised in the faculty facilities, were used at embryonic day 8 (E8) as previously described ^13,21^. Eggs were incubated at 37°C with 60% humid atmosphere (model 80/AD, Yonar SRL, Buenos Aires, Argentina). Chicken embryos were sacrificed by decapitation. All experiments were performed in accordance with the Use of Animals in Ophthalmic and Vision Research of ARVO, approved by CICUAL (Institutional Committee for the Care and Use of Experimental Animals, School of Chemistry, National University of Cordoba; RD-2025-1475-E-UNC-DEC#FCQ).

### Primary Cultures of Müller Glial Cells

Primary cultures of MC were purified from E8 chick retinas. Briefly, tissue was dissected in Ca+2-Mg+2 free Tyrode’s buffer (CMF) as previously reported ^13,14^. Briefly, cells were treated with papain (50 µg/ml) for 20 min at 37°C and after dissociation, the cells in suspension were seeded in Petri dishes and grown in DMEM supplemented with Fetal Bovine Serum (FBS) 10% for one week. Cell cultures were then trypsinized and re-plated in petri dishes or multiwell plates depending on the experiment and maintained for another week with FBS 10%, when we obtain purified glial cultures, virtually free of neurons. Primary cell cultures were previously characterized to determine the glial identity of cells ^14^. Results showed that most cells in the cultures co-express Glial Fibrillary Acidic Protein (GFAP) together with Glutamine Synthetase (GS) or Glutamate Aspartate Transporter 1 (GLAST1), strongly indicating that they are retinal MC. Cultures were incubated at 37°C under constant 5% CO2-air flow in a humid atmosphere (MCO 175, SANYO) across the whole experiment.

### Chick embryo retinal explants

Chicken embryos were sacrificed by decapitation at embryonic day 8 (E8) as previously described ^21^. After enucleation, eyes were hemisected equatorially and the gel vitreous was removed from the posterior eyecup. Tissue was embedded in DMEM for 2h in darkness and maintained in 5% CO2 at 37 °C. Afterwards, illumination protocols were carried out as indicated in the figures.

### Human MIO-M1 cell culture

The human Müller cell line (Moorfields/Institute of Ophthalmology-Müller 1) MIO-M1, was generated and previously characterized by G. Astrid Limb lab ^22^ and kindly provided by Dr. Cecilia Sanchez (CIBICI-CONICET-UNC). Cells were cultured in 24/96-well plates (Biologix), with 5,000/1,000 cells seeded per well respectively, using modified DMEM supplemented with 10% FBS and penicillin/streptomycin, and maintained in 5% CO2 at 37 °C.

### HEK-293 cell culture

Human embryonic kidney 293 cells (HEK 293T/17, ATCC) were seeded (15,000/plate) and cultured using modified DMEM supplemented with 10% FBS and penicillin/streptomycin, and maintained in 5% CO2 at 37 °C.

### Experimental procedures

In primary cultures at 80% cell confluency culture media was replaced with DMEM without FBS for 40 min. Cells were then exposed to BL (470–490 nm, peak at 480 nm LED, 1400Lx) and samples were collected at the indicated time points in the corresponding figure legend. In another set of experiments cell cultures were incubated with the retinal bleacher hydroxylamine (30 mM, 40min in DMEM, Taurus), the calcium inhibitor Thapsigargin (TG 2 μM), or DMEM supplemented with 9-*cis* retinal (0.6mM); illuminated with red light (630–650 nm, peak at 640 nm LED, 48 μW/cm2); or stimulated with forskolin (10/100μM, Caiman) and samples collected at the indicated time points in the corresponding figure legend. Similar conditions were used for sampling HEK-293 cells and retinal explants.

For proliferation and differentiation evaluation cell cycle synchronization at G0/G1 phase was achieved by serum-starvation for 24h at 37°C in a humidified 5% CO2 atmosphere. Afterwards cells were relieved into cell cycle by adding 10%/20% FBS; illuminated with BL and maintained in different media conditions as indicated in the corresponding figure.

### cAMP radiobinding-protein (RBP) assay

Following treatment, each well (icAMP) and well supernatant (e-cAMP) was extracted with 1min incubation with 95%v/v ethanol. In all cases, no PDE inhibitors were used during cAMP accumulation assays. Extracts were then evaporated and residues re-suspended in RBP-buffer (50mM Tris-HCl, 4mM EDTA, pH 7.4, 0.1% BSA). Samples were labeled with consecutive numbers and the corresponding measurements were performed blinded to the experimental group. cAMP content was determined by a competitive RBP assay for PKA using [3 H]-cAMP ^23^. Titrated PKA was incubated in equilibrium conditions (2h, 4°C) with different samples or cAMP standards (0.1-90pmol) in the presence of 2nM [3 H]-cAMP (20.7 Ci/mmol, PerkinElmer, #NET1161250UC). The bound fraction was separated by carbon-dextran precipitation followed by centrifugation (2000g, 15min, 4°C), and Optiphase HiSafe3 scintillation cocktail (PerkinElmer) was added to each supernatant for counting in a Pharmacia Wallac 1410 counter. Sample cAMP concentrations were determined by interpolating from the displacement curves obtained from cAMP standards using Prism 8.0 (GraphPad Software). No differences were observed for e-cAMP levels (data not shown), the result section describes the observed effects for i-cAMP.

### Immunocytochemistry (ICC)

Cultured cells were fixed for 15 min in 4% paraformaldehyde in phosphate-buffered saline (PBS), followed by methanol incubation for 10 min and washed in PBS. Afterwards, samples were treated with blocking buffer (3% bovine serum albumin, 0.1% Tween 20, 1% glycine, 0.02% sodium azide in PBS) and incubated overnight with the primary antibodies for: p-CREB; Vimentin; SOX2, RGR, GS,Tbr2, Brn3, GFAP, p53, or G3BP1 (See Suppl. Table 1). They were then rinsed in PBS and incubated with the secondary antibodies (1:1,000) for 1 h at RT. Samples were incubated with DAPI (3 μM). Coverslips were finally washed thoroughly and visualized by confocal microscopy (FV1200; Olympus,Tokyo, Japan) ^13^. Microscopy images were acquired at CEMINCO (Centro de Micro y Nanoscopía de Córdoba, Argentina, https://ceminco.conicet.unc.edu.ar/)

### Image Processing

For measuring the number of cells with positive staining microscopic photographs were acquired with an Olympus camera in a Fv-1200 confocal microscope. Analyzed images for all the groups were captured at the same magnification (40X), microscope setting, and stored as “oib” files for immunohistochemistry signal analysis. Each file was imported by Image J software (Image J 1.8.0 bundled with 64-bit Java, NIH, USA). The background signal was measured in negative controls and subtracted in all channels, then a threshold algorithm was used for the different markers to generate binary masks and selections of immunoreactive cells was completed with nuclear counting with DAPI staining. All imaging process steps were performed using specific Macros to avoid the operator factor.

### Calcium Imaging

Cells were grown in an 8-well Lab-Tek recording chamber (NuncTM, NY-USA). On the day of the experiment, MCs were incubated with 0.1% of pluronic acid F-127 and 5 μM of the Ca2+indicator dye (Calcium Orange/AM, Invitrogene), for 1 h at 37°C, under darkness condition. The cells were washed three times with CMF and then Ca2 +was measured in DMEM by exciting the indicator at 543 nm for 1 min. Cells were then stimulated with a BL pulse (20 s) ^13,14^ , and imaged for another 5–8 min. In order to interfere with light-driven Ca+2 response in MC, Thapsigargin (TG 2 μM) was added to the medium for at least 20 min before commencing the recordings. The fluorescence imaging technique was performed by confocal microscopy with an Olympus FluoView-1,000 microscope. The Ca2+fluorescence intensity ratio (F/F0) was calculated as a function of time and total average changes in fluorescence intensities along the whole experiment after the stimulus and are expressed as ΔF to compare between experimental groups. Microscopy images were acquired at CEMINCO (Centro de Micro y Nanoscopía de Córdoba, Argentina, https://ceminco.conicet.unc.edu.ar/)

### Retinal fixation and sectioning and Immunohistochemistry

Samples were fixed for 30 min at 20°C in 4% paraformaldehyde with 3% sucrose in 0.1 M phosphate-buffered saline (PBS), pH 7.4. Fixed samples were cryo-protected in PBS plus 30% sucrose, soaked in embedding medium (O.C.T. compound, Tissue-Tek) for 10 min and freeze-mounted onto aluminum sectioning blocks. Transverse sections nominally 14 µm thick were cut consistently and thaw-mounted on glass slides. Sections were air-dried and stored at −20°C until use. Immunohistochemical detection was performed following the same steps as for ICC. Microscopy images were acquired at CEMINCO (Centro de Micro y Nanoscopía de Córdoba, Argentina, https://ceminco.conicet.unc.edu.ar/)

### RNA Isolation and Polymerase Chain Reaction

Total RNA from MC cultures was extracted using the TRIzolTM kit for RNA isolation (Invitrogen). RNA integrity was checked and quantified by UV spectrophotometry (Epoch Microplate Spectrophotometer, Biotek). Finally, 2 mg of total RNA was treated with DNAse (Promega) to eliminate contaminating genomic DNA. cDNA was synthesized with M-MLV (Promega) using Random Primers (Promega) as previously described ^13^. The oligonucleotide sequences used for PCR from the *G. gallus* sequences were as follows: <u>PAX6:</u> Forward GGGAGTAGAGGCACGCAGATGT Reverse GGAGTCGCTACTCTCGGCTTACTA

TATA-binding protein (TBP):

Forward CCGGAATCATGGATCAGAACA

Reverse CGGCATCATCGGGCTAAA

For Polymerase Chain Reaction initial denaturation step was set for 1 min at 94°C, 35 cycles of 60 s at 94°C, 50 s at 60– 65°C, 90 s at 72°C, and a final 5 min elongation step at 72° C. Amplification products were separated by 2% agarose gel electrophoresis and visualized by ethidium bromide or sybersafe (InvitrogenTM) staining.

### Cell Cycle Progression Analysis

MCs cells were arrested in serum-free DMEM for 24 h. In a first set of experiments, cell cycle length in MC primary cultures was identified by stimulation to re-enter the cell cycle with 20% FBS and samples were taken at 0, 6 and 10h later. In another set MC primary cultures were illuminated at the beginning of the cell cycle and transferred to 10%FBS medium, sampling was performed 10h later. Cells harvested by trypsinization were washed in cold phosphate buffered saline (PBS) and fixed with cold 70% ethanol at 20°C for at least 24 h. Cell pellets were resuspended in 150 µL of staining solution (PBS containing 50 µg/mL propidium iodide and 10 µg RNAse A) as reported ^24^. Cell cycle analysis was performed by the flow cytometer BD LSRFortessa and FlowJo software (Verity Software House, Topsham, ME, USA).

### Cell Migration Analysis

Cell migration was analyzed using the wound healing assay. Briefly, in a 24-well plate, a “wound” was created on the cell monolayer (t=0), and afterwards cells were either stimulated with blue light or kept in the dark. Images were captured with a Leica DMI 8 at t=0 and t=24h, and the percentage of wound closure was calculated. ImageJ software was used for image processing.

### Cell Proliferation Rate Analysis

Cell proliferation in culture was evaluated using the resazurin assay, following the manufacturer’s protocol. Briefly, cells were seeded in a 96-well plate and cell-cycle arrested for 24h in serum-free culture medium. Subsequently, cells were exposed to blue light and stimulated to re-enter the cell cycle by adding DMEM supplemented with 0, 10 or 20% FBS and 10% v/v alamar Blue dye (Invitrogen). Following 4 hours of incubation, fluorescence was measured (t=0) using a BioTek microplate reader. BL stimulation was sustained at the beginning of each cell cycle (every 12h) and fluorescence for each condition was recorded every 24 hours over 4 days and referenced to time zero (t=0).

### Western blotting

Homogenates primary cultures were resuspended in RIPA buffer (50 mM Trise HCl, pH 8.0, with 150mM sodium chloride, 1.0% triton, 0.5% sodiumdeoxycholate, and 0.1% sodium dodecyl sulfate) containing protease inhibitors (Sigma Aldrich) and processed for WB. Homogenates were resuspended in sample buffer and separated by SDS-gel electrophoresis on 12% polyacrylamide gels (50 mg total protein/lane), transferred onto nitrocellulose membranes, blocked for 1 h at RT with 5% Bovine Serum Albumin (BSA) in PBS and then incubated overnight at 4°C with specific antibodies for RGR, GS, tubulin, III-β-Tubulin; β-actin; and GFAP. Membranes were washed three times for 15 min each in washing buffer (0.1% Tween 20 in PBS) and incubated with the corresponding secondary antibody in the incubation buffer during 1 h at RT, followed by three washes. Membranes were scanned using an Odyssey IR Imager (LI-COR Biosciences).

### Statistics

Statistical analyzes involved t-test, one-way/two way/repeated meassures analysis of variance (ANOVA) followed by Bonferroni post-hoc comparison when appropriate. Otherwise, Mann-Withney (M-W) or Kruskal-Wallis (K-W) tests were used when normal distribution or homogeneity of residuals was infringed with pairwise comparisons performed by the Dunn’s test when appropriate. In all cases significance was considered at p < 0.05.

## Results

### Blue light triggers cAMP signalling in Müller glial cells

Our previous observations demonstrated the intrinsic photosensitivity of MCs, evidenced by calcium release from internal stores in response to BL ^13,14^. Although most cells in primary MC cultures express non-visual opsins, only a subpopulation (40–55% of analyzed cells) responds to light with intracellular calcium increases ^13,14^. To assess whether an alternative signaling pathway could be triggered by the blue-sensitive opsin in MCs, we evaluated cyclic nucleotides content in MCs cultures after BL by RBP determination of cAMP levels, as described in methods. A 1-min BL stimulus induced a significant increase in intracellular cAMP, peaking at 10 min post-stimulus and returning to basal levels by 30 min (H_(3)_=12.13; p=0.0069 . Fig. 1A). Further determinations at 10 min post-stimulus confirmed a significant increase in cAMP levels following 1-min BL stimulation (H_(4)_=19.57; p=0.0006, dunn’s test: p<0.05 compared to Dark conditions. Fig. 1B) This response was blunted by prior incubation with the non-specific opsin antagonist hydroxylamine (Dunn’s test: p>0.05 compared to Dark conditions. Fig. 1B). Given that hydroxylamine effectively blocks opsin-mediated calcium increases in both neurons and MCs ^12,13^; our results indicate that BL activates a non-visual opsin to drive cAMP signaling in MCs. Consistent with the previously reported wavelength sensitivity of calcium responses ^14^, this pathway is wavelength-specific; no increases in cAMP were detected 10 min after red light stimulation (1 min, RL; dunn’s test: p>0.05 compared to Dark conditions. Fig. 1B). As a positive control, cAMP levels were elevated using forskolin (FSK), a direct adenylyl cyclase activator ^25^. FSK treatment (10μM/100μM for 5 min) promoted a pronounced increase in cAMP levels in MC cultures at 10 min (FSK, H_(2)_=10.03; p=0.006; dunn’s test: p<0.05 compared to basal conditions. Fig. 1C).

**Figure 1.**
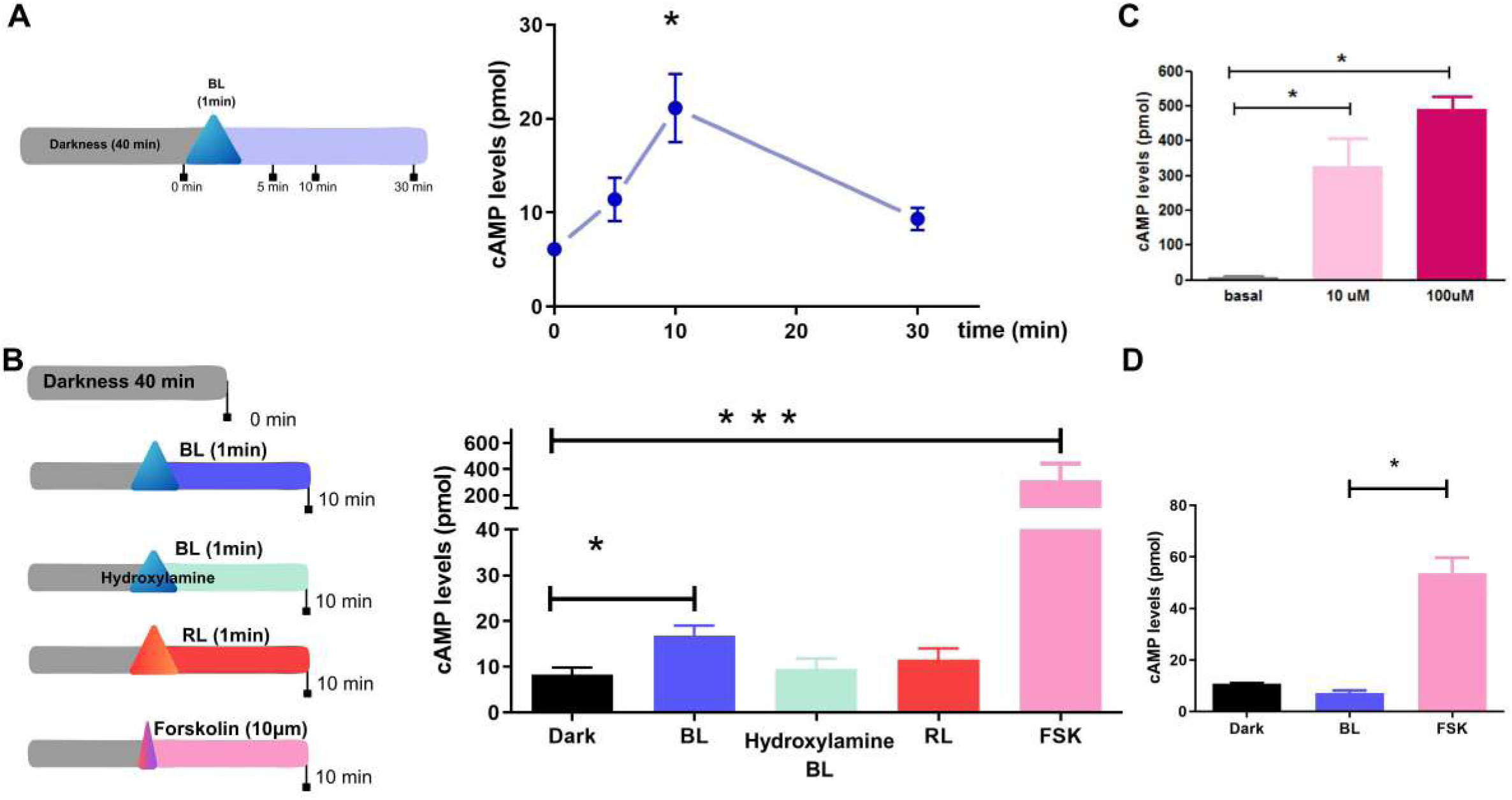
Blue light triggers cAMP responses in Müller glial Cells. The experimental protocol for evaluating cAMP levels after a blue light pulse (1 min, BL) is shown in left scheme (A). The graph shows the time line for cAMP intracellular levels in MC primary cultures; highest levels were observed by 10min after the BL stimulus (A). n= 5. Experimental protocols for evaluating cAMP levels by 10 minutes under different experimental conditions is shown in left panel (B). The graph shows a positive response in cAMP levels after the BL stimulus and Forskolin stimulation (FSK); previous incubation with the retinal bleacher Hydroxylamine blunted the cAMP response by BL, whereas red light stimulation (RL) failed to induce the second messenger, n=4-13. Positive controls assessing cAMP levels elicited by different FSK concentration in MC (C). Cell type specificity was addressed by cAMP levels evaluation in HEK-293 cells in basal conditions, after a BL pulse or FSK stimulation (D); n=3. The bars represent the mean±SEM, *p<0.05 and ***p<0.001 different from Dark/basal levels.

Additional control was carried out to assess the specificity of the observed photic responses, by assessing cAMP levels in HEK-293 cells (Fig. 1D). HEK-293 cell line does not express Opn3 nor Opn5 ^13,26,27^ and does not show calcium responses to BL stimulation^28^. In the same way, no significant changes are detected in cAMP levels of HEK-293 cells 10 min after BL (1 min; H_(2)_= 7.2; p=0.027; Dunns test: p>0.05. Fig. 1D), still they do respond to FSK stimulation (10μM for 5 min; Dunns test: p<0.05. Fig. 1D).

### CREB activation in Müller glial cells by Blue Light *in vitro* and *ex vivo*

Amid the various downstream effectors for cAMP, activation of the protein kinase A (PKA) and consequently phosphorylation of the Ser133 residue in cAMP-responsive element binding protein (p-CREB) drives the transcription of various target genes ^29^. Hence, we next examined the temporal course of CREB activation by BL in two experimental models. *In vitro* experiments included immunolabeling of p-CREB in MCs primary cultures. The percentage of p-CREB positive nuclei indicate a time-dependent activation of CREB by sustained BL stimulation (15-30-60 min. Fig. 2 and Fig. 3. H_(3)_= 41.72; p<0.0001). Particularly, significant activation is achieved after 30 min of BL (Dunns test p<0.0001 different to DARK controls. Fig. 2A) and reaches higher levels by 60 min of BL as compared to DARK conditions (Dunns test p<0.0001. Fig. 2A).

**Figure 2.**
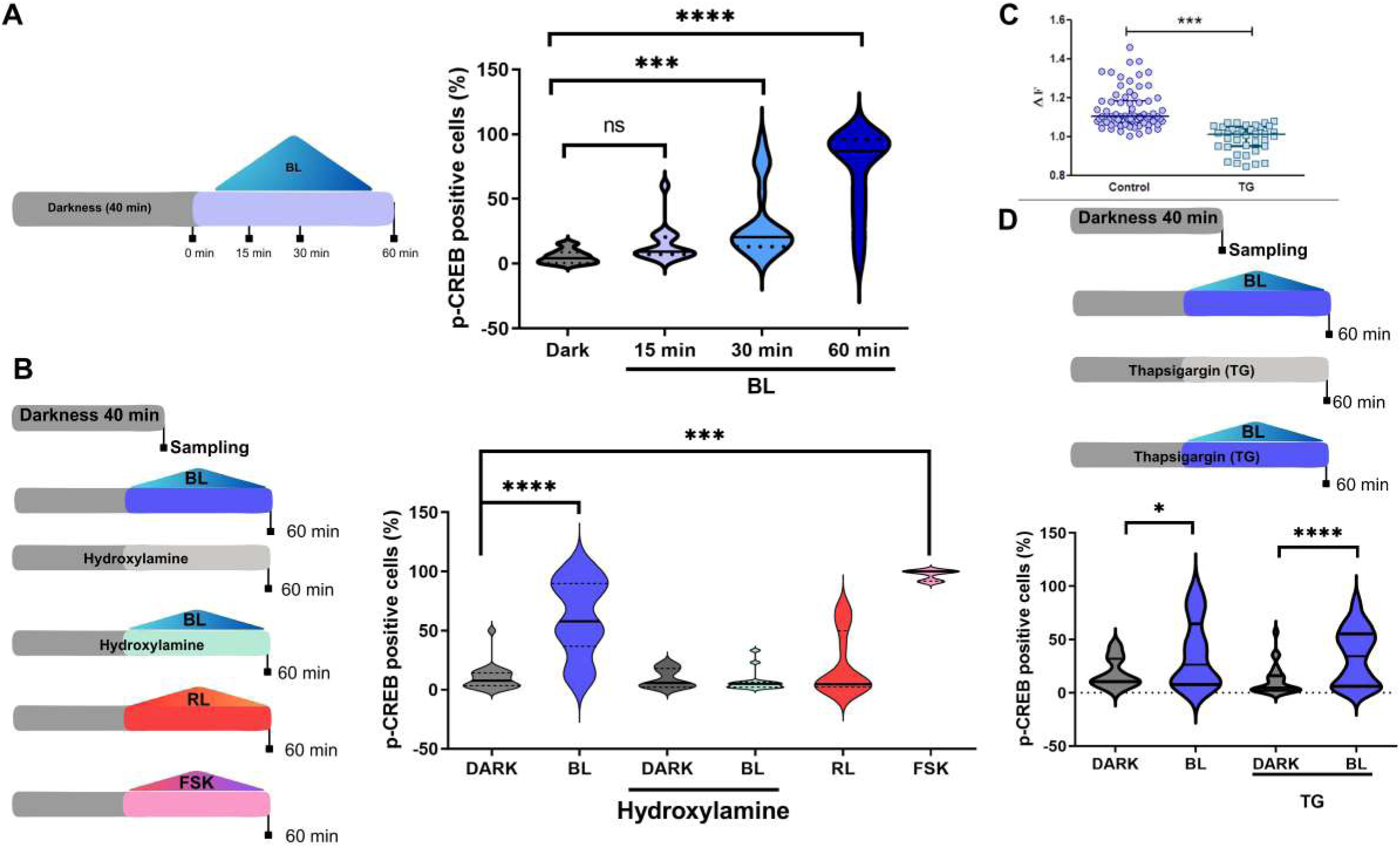
**Temporal course of CREB activation by Blue Light**. Experimental protocol for evaluating CREB activation after blue light (BL) is shown in left panel (A). The graphs shows the percentage of p-CREB positive nuclei in MC primary cultures, showing a time-dependent activation of CREB by sustained BL stimulation by 15, 30 and 60 min (A). Significant activation is achieved after 30 and 60 min BL as compared to Dark controls. n = 5 independent experiments. The schemes in left panel (B) shows the different experimental conditions for evaluating CREB activation by 60 min after stimulation. The BL-driven CREB activation depends on opsin-mediated signalling (as hydroxylamine prevented p-CREB induction) and is wavelength specific (no activation observed with red light, RL). Significant p-CREB levels are achieved by Forskolin stimulation (FSK) (B). Pharmacologic depletion of calcium internal stores with Tapsigargin (TG) effectively blocks calcium signalling elicited by BL in MC cultures (C, graph showing the median with interquartile range). When administered for p-CREB evaluation (scheme in D) it does not modify blue-light driven CREB activation (D). The p-CREB graphs show the median and interquartile range. *p<0.05 and ***p<0.001 ****p<0.0001different from basal/ Dark levels.

**Figure 3.**
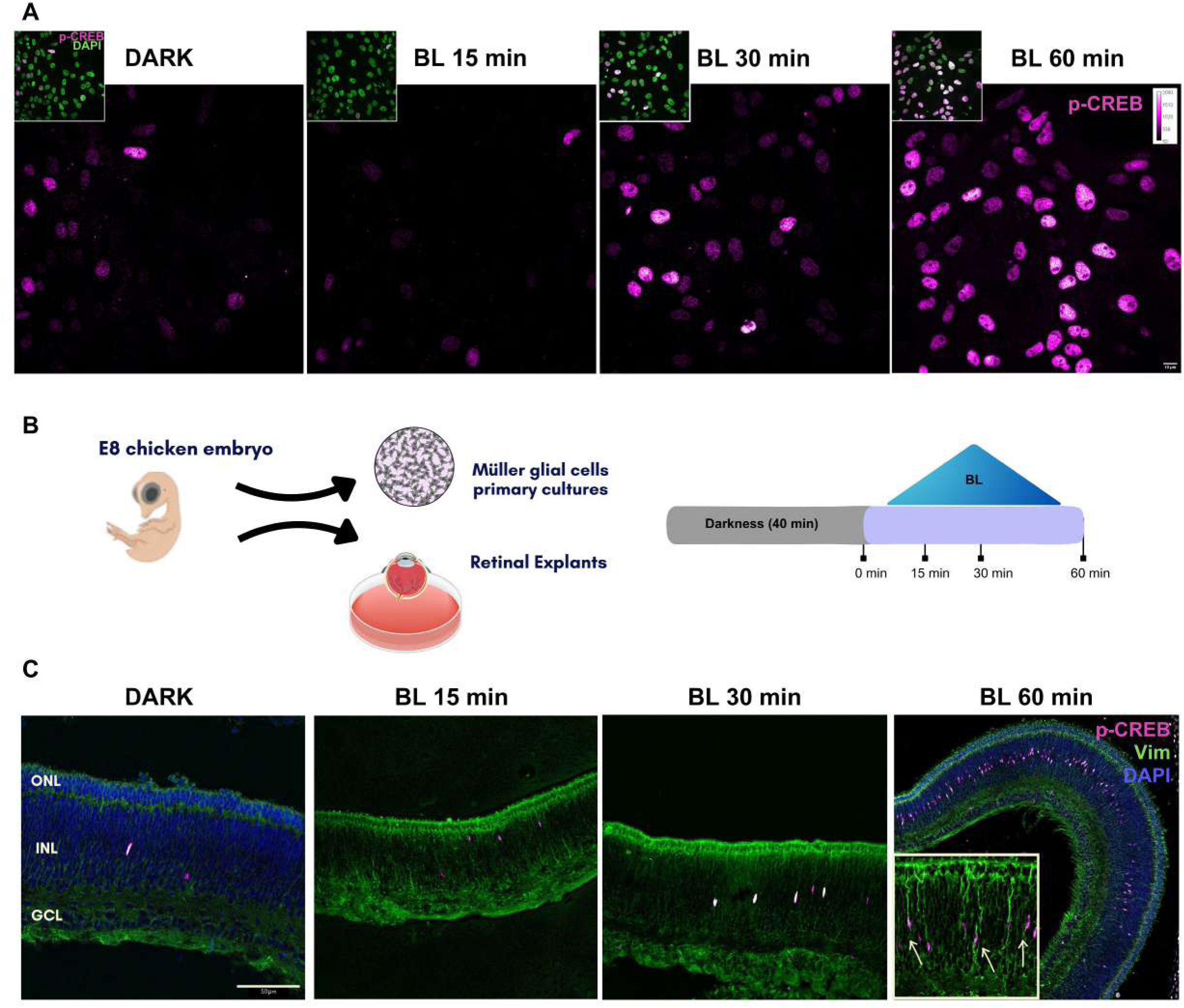
**Temporal course of CREB activation in Müller glial cells by Blue Light *in vitro* and *ex vivo***. Upper panel (A) shows representative microphotographs of Dark and BL stimulated MC in primary cultures and quantified in figure 2. p-CREB staining is shown in magenta gradient and the total number of cells present is observed by merge with Dapi staining (in green) shown in the inset for each analyzed time-point. The experimental protocol was conducted in two different models as illustrated in the central scheme (B) for evaluating MC activation *in vitro* and *ex vivo* conditions. BL stimulation to dark-adapted E8 retinas increases p-CREB expression at different time points (C, by 15, 30 and 60 min; n=3 independent experiments). Interestingly, this activation is observed in regional patches along the inner retina and is mainly restricted to the inner nuclear layer where MCS nuclei are located. In fact, p-CREB immunostaining (magenta) colocalizes with the glial marker Vimentin (green) as indicated by the arrows in the inset at 60 min (C). ONL: Outer Nuclear Layer; INL: Inner Nuclear Layer; GCL: Ganglion Cell Layer

Consistent with cAMP dynamics, 1 h of BL stimulation drives CREB activation through an opsin-mediated pathway, an effect blocked by prior incubation with hydroxylamine (H_(5)_= 48.40; p<0.0001. Fig. 2B), whilst it did not modify basal levels of p-CREB in darkness conditions (Dunns test: p>0.05 for DARK vs DARK with Hydroxylamine. Fig. 2B). In the same direction, wavelength sensitivity was corroborated after 1h of red light stimulation as MCs do not achieve any significant activation of CREB (RL, p>0.05 compared to DARK. Fig. 2B). Positive control with FSK (10μM, 30 min) strongly drives CREB activation reaching highest values of p-CREB positive nuclei at the evaluated time point (FSK. Fig. 2B).

Considering our previous observation of BL-induced calcium release from internal stores in MCs ^14^, we further tested a possible overlap for calcium and cAMP signalling in the BL-driven CREB activation. Pharmacologic depletion of calcium internal stores with thapsigargin effectively decreases the overall calcium responses in MC cultures (Mann Withney U= 156; p<0.0001. Fig 2C); whilst it does not affect BL-induction of p-CREB with a significantly increased percentage of p-CREB positive nuclei than controls kept in darkness (H(3)= 21.98; p<0.0001. Fig. 2D). Indeed, the overall levels of CREB activation was not affected by thapsigargin preincubation, with similar levels between DARK and BL at the different conditions (DARK vs TG+DARK , and BL vs TG+BL, p>0.05).

In order to extend our observations we tested for CREB activation in retinal explants, an *ex-vivo* model that allows us to elucidate cellular responses in the complex retinal network, maintaining tissue integrity and cell-cell connectivity^21^ (Fig. 3B). BL stimulation to dark-adapted E8 retinas temporarily increases p-CREB expression (Fig. 3C), resembling the phenomena observed in cultured MCs (Fig. 3A). Interestingly, this activation is observed in regional patches along the inner retina and is mainly restricted to the inner nuclear layer (Fig. 3C), where MC nuclei are located. In fact, p-CREB immunostaining colocalizes with the glial marker Vimentin (Fig. 3C). This p-CREB/vimentin positive MC immunostaining pattern resembles the already identified opsin-expressing MCs in the embryonic retina ^28^. Notably, pharmacological induction of p-CREB by FSK treatment (10μM, 30 min) in retinal explants, mirrors the glial activation pattern observed upon 30 min of BL stimulation (Suppl. Fig. 1A). Previous incubation with the chromophore 9*-cis* retinal in dark-adapted retinas causes an overall p-CREB activation after 1h of BL, observed as p-CREB positive nuclei in all of the nuclear cell layers at the developing retina (Suppl. Fig. 1B).

### Proliferative capacity in Müller glial cells after BL stimulation

The nuclear activated CREB, forming homodimers or heterodimers, binds to the cAMP response element (CRE) in the promoter region of target genes governing cell differentiation, proliferation, migration, apoptosis, metabolism, and more ^29^. To a better understanding of BL-triggered cellular responses, we first determined cell cycle progression in MC cultures kept in darkness conditions under proliferative conditions (20% FBS), which indicates a 12h cell cycle for MCs undergoing cell division (Suppl.Fig. 2).

To assess how BL stimulation influences cell cycle progression, we arrested MC cultures for 24 h using serum-free DMEM (0% FBS). We then exposed the cells to BL for 1 h at the beginning of the cell cycle (keeping controls in the dark) before releasing them into the cell cycle with 10–20% FBS. Flow cytometry was performed 10 h after BL stimulation to evaluate cell cycle progression (Fig. 4A). At this time point, BL-stimulated MC cultures show significant different cells percentage at different cell cycle stages as compared to DARK controls maintained in proliferative conditions with 20%FBS (Two way ANOVA-RM Interaction cell cycle stage*group F_(4, 64)_=3.65, p=0.0097). When analyzed in further detail this discrepancy is in fact reflecting concomitant increased cell percentage at S phase and decreased percentage in G2/M phase in BL-exposed cultures (at both stages BL 10%FBS vs DARK 20%FBS p<0.05 by Tuckey *post hoc* test), thus suggesting that proliferative capacity of MCs is compromised after BL stimulation by modifying cell cycle timing (Fig. 4B). The effects of BL were further evaluated considering MC migration and proliferation by analyzing the results after 1h-BL exposure at the beginning of each cell cycle (Scheme in Fig 4D). Strikingly, BL exerts an inhibitory effect over migratory and proliferative capacity of MCs (maintained in 10% FBS media), observed as lower wound closure percentage by 24h (t_(141)_=2.41, p<0.05. Fig. 4C) and diminished growth rate monitored along 4 days (Two way ANOVA RM interaction time*groups F_(3,282)_=7.95, p<0.0001; significant differences at all time points by Bonferroni *post hoc* test. Fig. 4F). Moreover, when tested at different serum concentrations substantial differences are observed on the inhibitory effect of BL over cell proliferation: while the growth rate is significantly lowered at 0% FBS conditions (Two way ANOVA RM interaction time*groups F_(3, 210)_= 212.5, p<0.0001; significant differences at all time points by Bonferroni *post hoc* test. Fig. 4E); MCs maintained with 20% SFB sustain similar growth rate as DARK controls, thus blunting the inhibitory effect of BL (Two way ANOVA RM interaction time*groups F_(3, 27)_= 0.27, p=0.84. Fig. 4G).

**Figure 4.**
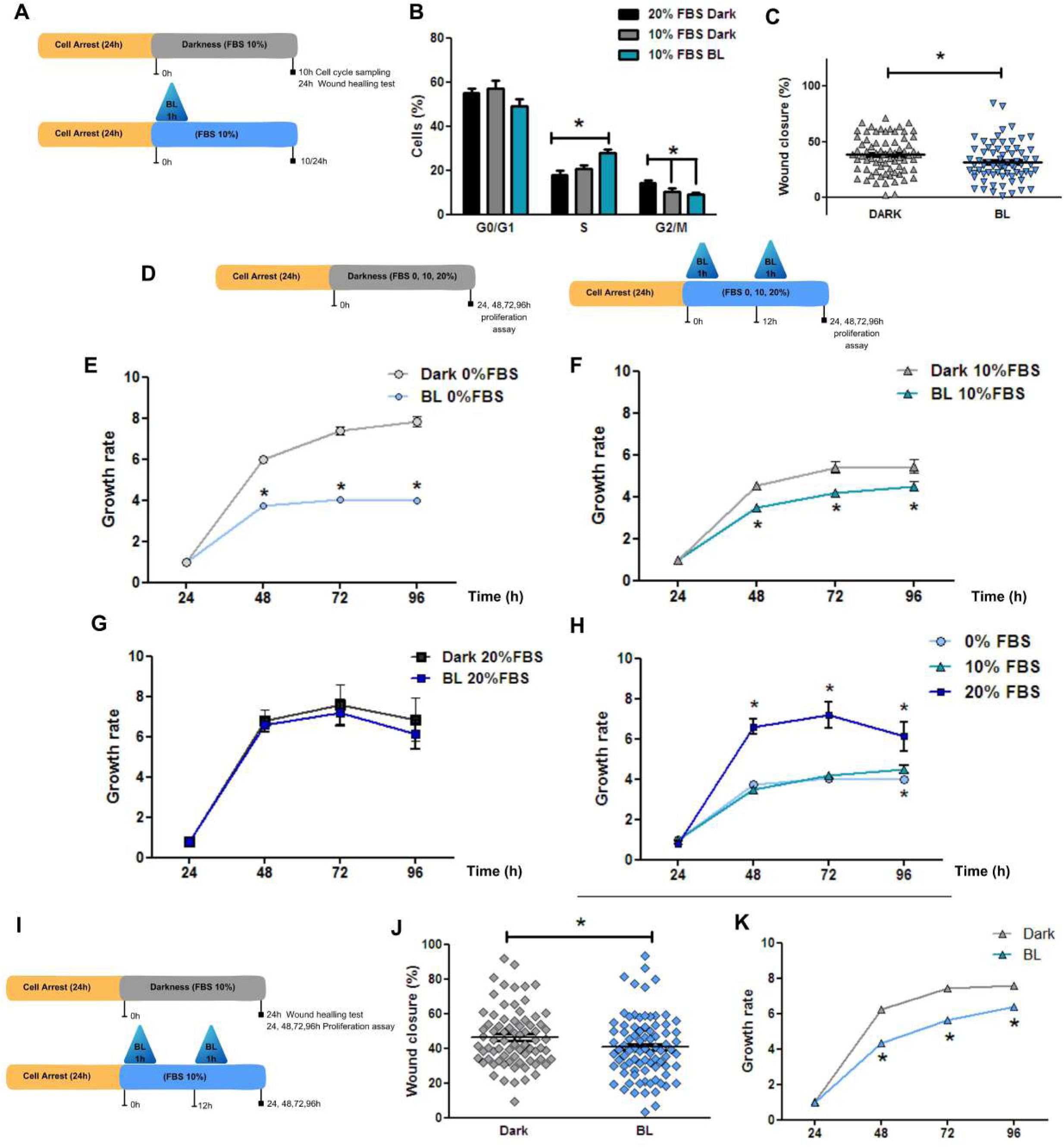
Blue light hampers Müller glial cells proliferation. The effects of BL on MC proliferation and migration were analyzed after 1h-BL exposure at the beginning of each cell cycle (schemes in A and D). BL increases the cell percentage of MC in the s-phase (B, n=11-13) and exerts an inhibitory effect over migratory capacity in differentiated MC (10% FBS culture media), observed as lower wound closure percentage by 24h (C, n=70-73 from 4 independent samples). MC proliferative capacity is hampered by BL stimulation in MC cultures (E-0% FBS, and F-10% FBS, respectively), observed as diminished growth rate monitored along 4 days(n=36 from 4 independent samples). No signifficant effect of BL stimulation over MC proliferations is achieved in proliferative conditions (G-20% FBS, ). Proliferative capacity is hampered to similar rates by BL stimulus when working with 0% or 10% FBS supplemented culture media (H). Under the same protocol (scheme in I) MIOM1 cells were evaluated after BL stimulus. Immortalized human MC stimulated with BL show significant decreased migration (J, n=75/83 from 2 independent experiments) and proliferative capacity (K, n=94/96).The graphs show the mean±SEM, *p<0.05 different from Dark conditions.

Notably, the effects of BL over migration and proliferation is also conserved in the human-derived Müller cell line MIO-M1, which are photosensitive cells ^30^. MIO-M1 cell cultures were submitted to the same experimental protocol (Fig.4I) and tested in the wound healing and proliferation (alamar blue) assays. 1h-BL exposure at the beginning of each cell cycle inhibits migratory and proliferative capacity of MCs (maintained in 10% FBS media), observed as lower wound closure percentage by 24h (t_(156)_=2.07, p<0.05. Fig. 4J) and diminished growth rate along 4 days (Two way ANOVA RM interaction time*groups F_(3,558)_=132.6, p<0.0001; significant differences at 48, 72 and 96h by Bonferroni *post hoc* test. Fig. 4K).

Given that the maximal effects of BL on proliferative capacity were observed in the absence of serum (0% FBS; Fig. 4E), and that BL stimulation reduced proliferation to comparable growth rates regardless of 10% FBS supplementation (two-way RM-ANOVA, interaction time*groups: *F*_(6, 261)_ = 25.57, *p* < 0.0001; Bonferroni post-hoc test, *p* > 0.05; Fig. 4H), all subsequent experiments were conducted using primary MC cultures maintained at 0% FBS.

### Blue light decreases Pluripotency signature in MCs

During retinal development, SOX2 expression clearly aligns with periods and areas of high retinal progenitor cell proliferation. Therefore the disappearance of SOX2 is a straightforward indicator of cellular maturation as progenitors finalize their neurogenic commitment. We next examined the temporal course of multipotency capacity in MCs by SOX2 immunolabeling at 6, 12 and 24 h after BL stimulation. The percentage of MCs expressing nuclear SOX2 is modified after BL stimulation, significantly decreasing by 12 and 24h (H_(3)_=32.03; p<0.0001; Dunns test: p<0.0001 for 12/24h vs DARK. Fig. 5A). In the same direction, *PAX6* transcript is decreased at the same time points after BL exposure (Fig. 5B), whereas stimulation with FSK (1µM) for 1h at the beginning of the cell cycle decreases SOX2 expression by 12h in MCs in cultures (Mann Whitney U= 179; p<0.001). So far, our results suggest that BL through cAMP signalling hampers proliferation and pluripotency capacity in MCs in cultures, possibly modifying their cellular fate.

**Figure 5.**
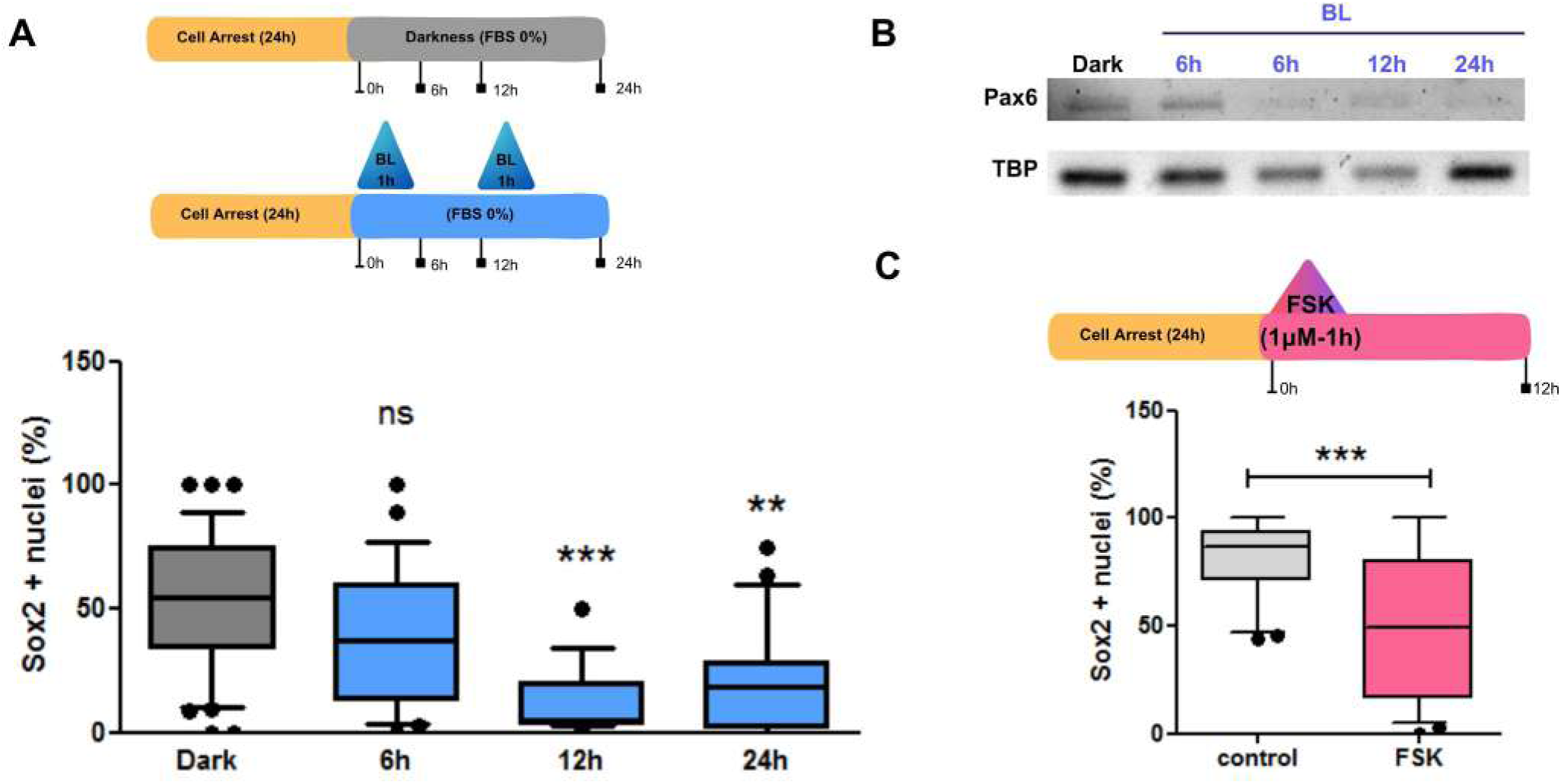
Blue light decreases pluripotency signature in Müller glial cells. Pluripotency of undifferentiated MC (0% FBS supplemented culture media) was assessed by Sox2 immunolabeling at different time points after 1h BL stimulation (Scheme in A). Decreased Sox2 positive nuclei are detected by 12 and 24h after BL stimulation (A), n=5 independent experiments. Lower levels of Pax6 transcript was also detected at the same time points (B). cAMP activation by Forskolin (FSK) stimulation promotes a similar decrease in Sox2 expression (C), n=5 independent samples. The graphs show the median and interquartile range, **p<0.01 and ***p<0.001 different from Dark/control levels.

### Blue light compromises MCs fate towards intermediate progenitors cells

In view of the last results, we checked for different cellular marker expressions in our experimental conditions, to identify the cellular identity of MCs cultures stimulated with BL. We evaluated the expression and distribution of the Müller glial marker RGR and the pan-glial marker GS 24 h post-stimulation using immunocytochemistry and western blot (Fig. 6). Immunolabeling revealed no changes in the cellular distribution of either marker (Fig. 6B, D). Similarly, protein expression levels remained unchanged for both RGR (Mann–Whitney *U* = 54, *p* = 0.48; Fig. 6C) and GS (*t*_(18)_ = 1.51, *p* = 0.15; Fig. 6E), indicating that BL exposure preserved Müller glial identity. We next examined whether BL triggers reactive gliosis or phototoxicity. Expression of the gliotic marker glial fibrillary acidic protein (GFAP) was unaltered in cellular distribution at 6 h post-stimulus (Fig. 6F) and in protein levels at 24 h post-stimulus (Mann–Whitney *U* = 21, *p* = 1.0; Fig.6E). To establish the safety threshold for BL exposure, we evaluated the pro-apoptotic marker p53 and the stress granule component G3BP1 under different light intensities (Suppl. Fig. 3A and B). At the experimental intensity of 1400 lx, no changes in p53 or G3BP1 immunostaining were observed. In contrast, exposure to 3000 lx induced nuclear translocation of p53 and stress granule formation. Together, these findings indicate that the responses observed under our experimental conditions reflect physiological BL-driven mechanisms rather than phototoxic stress or gliosis.

**Figure 6.**
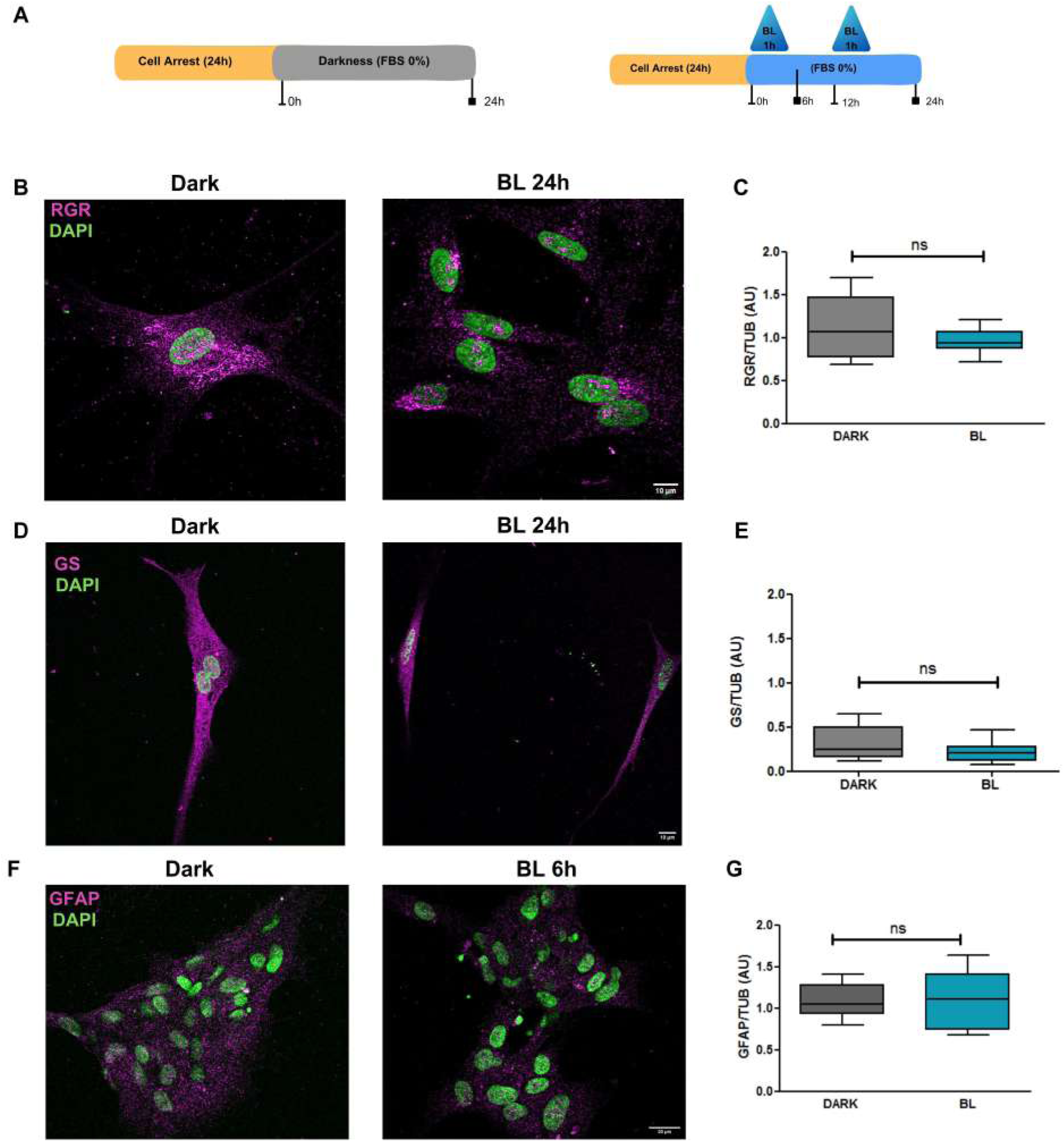
**Blue light does not modify Müller glial cells markers**. As shown in the scheme (A), the expression of glial markers was assessed 24h after BL stimulation in undifferentiated culture conditions (0% FBS). Immunostaining of retinal G protein-coupled receptor (RGR, in magenta at B panel microphotographs) and Glutamine Synthase (GS, in magenta at D panel microphotographs) showed no differences in location nor expression between dark and BL stimulated MC in culture. When analyzed by western blot, the levels of RGR (C, n=11) and GS (D, n=11) remained unchanged 24h after BL stimulations. Gliotic response by Glial Fibrillary Acidic Protein (GFAP) expression was evaluated short and long times after the stimulus (F and G). No difference in immunostaining was detected in MC cultures 6h after the BL stimulus (F), whereas no significant differences in GFAP levels are detected 24 after BL exposure (B, n=6-7). The graphs show the median and interquartile range.

Along development transcriptional factors are sequentially expressed to shape the ability of neural progenitors to proliferate and generate specific cell types. In particular, we evaluated staining of Tbr2 protein (gene: Eomes), a critical T-box brain transcription factor 2 known as the hallmark marker for intermediate progenitor cells (IPCs) in the developing central nervous system, which is mainly expressed in basal layers of the developing chicken retina at the E8 developmental stage (Fig. 7A). We examined Tbr2 expression in our MC cultures (scheme in Fig.7B) and evidenced an increase in Tbr2 immunostaining 48 and 72h after BL exposure compared to MCs maintained in darkness conditions (Fig. 7C and D), indicating that BL-stimulation in MCs could drive their cellular commitment towards neurogenic progenitors.

**Figure 7.**
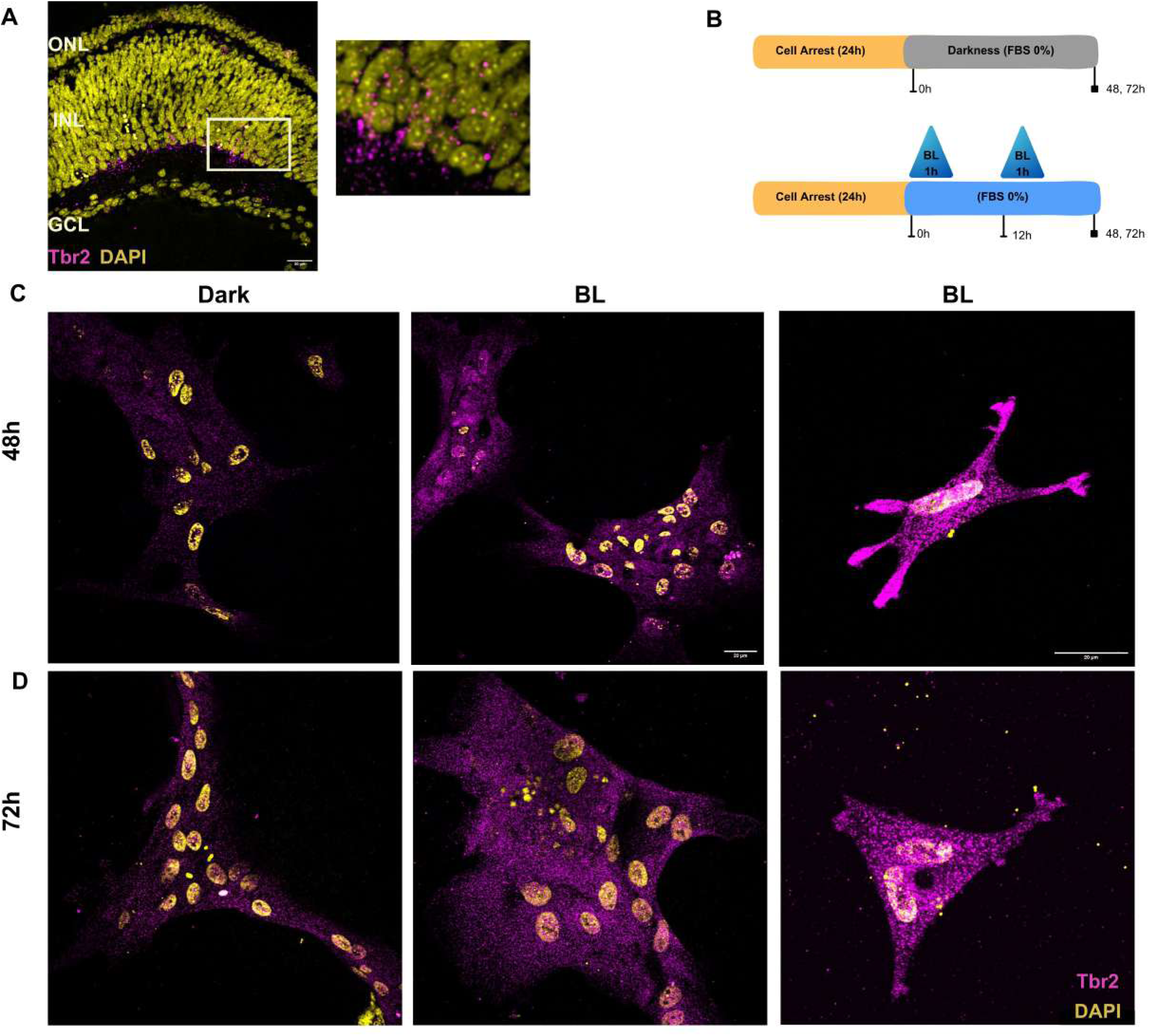
Blue light increases neuronal specification markers. Expression of the transcriptional factor Tbr2 (gene: Eomes, in magenta) was determined in E8 chicken retinas in basal conditions to confirm positive immunolabeling of the transcriptional factor in developmental stages. The selected area is zoom-in at the right microphotograph. ONL: Outer Nuclear Layer; INL: Inner Nuclear Layer; GCL: Ganglion Cell Layer. Expression of Tbr2 was assesed 48 and 72h after BL stimulation in MC primary cultures (scheme in B). The microphotographs are representative of increased Tbr2 immunostaining detected 48 (C) and 72h (D) in BL stimulated MC cultures.

### Blue light drives MCs differentiation into retinal neurons profile

Finally, to assess whether these MCs-derived neuronal progenitors could effectively differentiate into neurons we evaluated expression of neuronal markers in our cell cultures. For this experiment we illuminated MCs for 1h at the beginning of two cell cycles (12 and 24 h) after arrest. Following light stimulus cell cultures were maintained under two different cell culture conditions for 96 h: MC media (DMEM supplemented with FBS 10%) or neuronal media (DMEM supplemented with B27) (Fig. 8A-samples collected at 120 h). The two way ANOVA (medium condition/lighting condition) identifies a significant increase in the neuronal marker ß3-Tubulin protein expression (by western blot) after light stimulus (Fig. 8B; F_(1,29)_= 6.29, p<0.05) together with a light-directed increase in the percentage of cells expressing the transcriptional factor Brn3 regardless of the medium condition (Fig. 8C; F_(1,36)_= 11.56, p<0.01).

**Figure 8.**
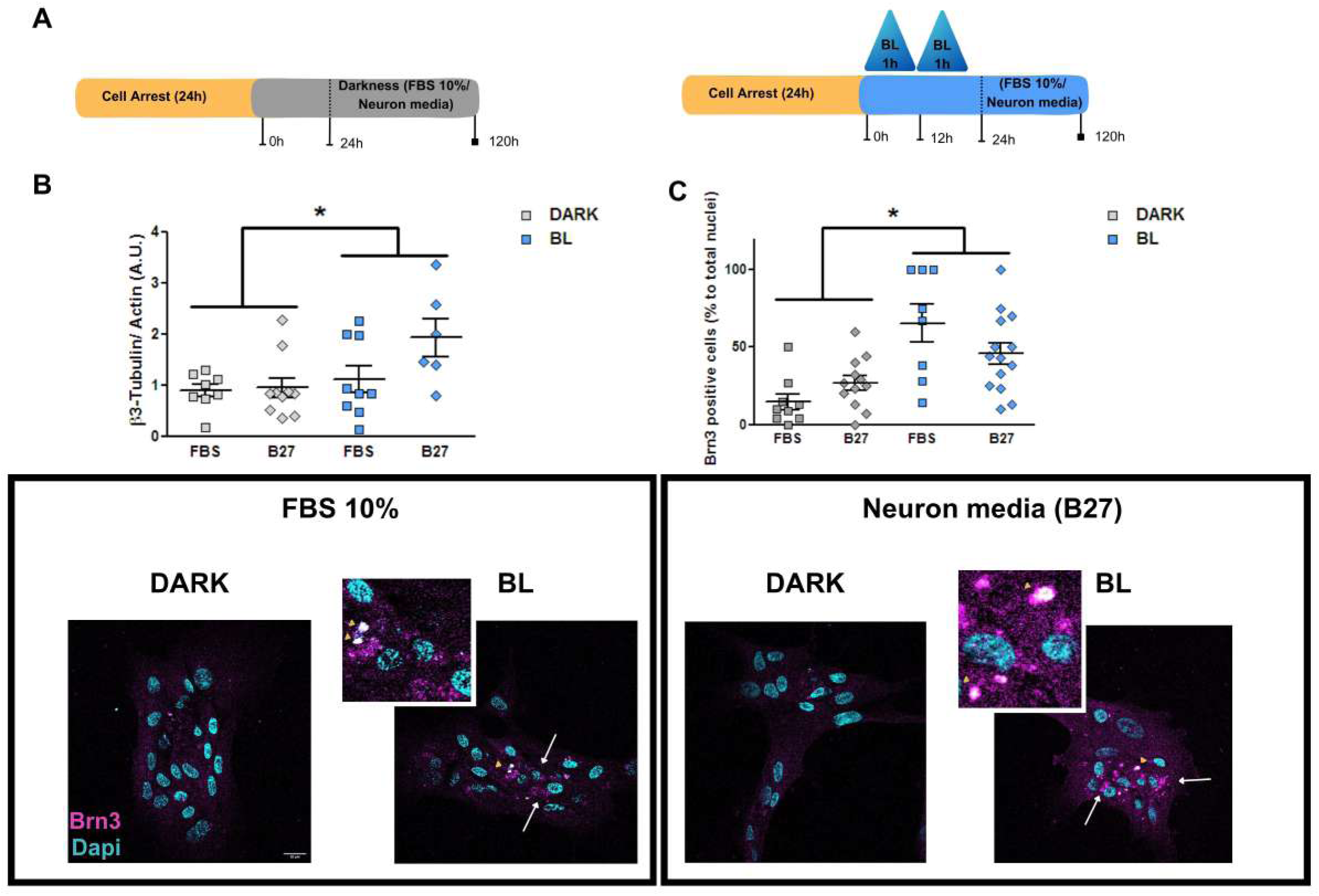
Blue light increases neuronal markers in MCs cultures. MC primary cultures were maintained and BL stimulated at the beginning of two cell cycles. Afterwards, cells were returned to darkness in two different medium conditions: MC medium (FBS 10%) or Neuron media (suplemmented with B27); and samples were taken 120h after the first light stimulation (Scheme in A). Following BL stiumlation a significant increase in the neuronal marker *β*3-tubulin (Tuj-1 clone) is detected (B, n=6-10) along with an increase in the number of positive cells expressing the ganglion cell specification marker Brn3, regardless the culture media conditions (C-Representative microphotographs on the bottom with zoom in at the yellow arrowhead. n=3 independent samples). The graphs show the mean±SEM, *p<0.05 different from Dark conditions.

## Discussion

Photoreception is an essential regulatory component in a plethora of physiological processes in ocular and extra-ocular tissues (i.e. skin, adipose tissue, blood vessels, brain) ^2,31–33^. In the last decades, non-visual opsins activation in ocular tissues has gained relevance in regulatory processes not related with image forming functions ^12,21,34–36^. We have recently described MC as a new photoreceptor cell within the embryonic retina expressing the non-visual opsins Opn3 and Opn5 and responding to BL pulses with calcium release from internal stores ^14,28^. In the present work, we extend the description of the signalling cascades triggered by BL and identify a light-directed cellular function in MC, unrevealing its potential in driving cellular fate.

We first characterized the time course of intracellular cAMP following a 1-minute blue light (BL) pulse, observing a peak at 10 minutes and a return to baseline by 30 minutes (Fig. 1).

This signaling cascade requires opsin activation and is wavelength-specific (Fig. 1), consistent with our previous findings regarding calcium responses ^14,28^. Furthermore, this BL-driven cAMP response depends on opsin expression, as it is absent in non-opsin-expressing HEK-293 cells^28^ (Fig. 1). Opsins constitute a monophyletic branch within the GPCR superfamily that lacks strict coupling to a single G protein across different cell types, tissues, and species^2,8,37^. Specifically, blue-light-sensitive Opn3 is phylogenetically linked to vertebrate visual opsins and teleost multiple tissue (TMT) opsin, both of which classically activate the *G_i_* pathway to decrease cAMP ^38^. However, experiments using Opn3 chimeras indicate that these opsins are also capable of elevating calcium and cAMP levels ^39,40^. Indeed, light-driven increases in both second messengers have been documented across multiple cell lines upon Opn3 or Opn4 stimulation ^6,41–47^.

Intracellular cAMP mediates cellular responses to external stimuli through downstream effectors ^48^. In the canonical pathway, cAMP activates protein kinase A (PKA) to regulate downstream targets, including the transcription factor CREB. Total CREB protein levels remain stable post-stimulation while the phosphorylated pool rapidly expands, indicating that signaling modulates CREB activity—specifically the magnitude and duration of nuclear p-CREB—rather than total protein abundance ^29,49^. Consistent with this mechanism, our results show time-dependent CREB phosphorylation in cultured MCs following BL stimulation, initiating at 30 minutes and peaking by 1 hour (Fig. 2A). CREB activation directly mirrors the preceding intracellular cAMP surge, similar to previous reports ^50^, and specifically, it shares core properties of a light-driven cellular response, including opsin dependence, wavelength specificity, and persistence after blocking intracellular calcium signals with thapsigargin (Fig. 2C, D). Most outstanding is the fact that we evidenced a similar p-CREB activation pattern in the embryonic tissue, with similar activation time-line and cellular specificity, as positive p-CREB are also positive for the glial marker vimentin in the inner nuclear layer of the chicken retina (Fig. 3C); supporting cAMP-mediated signaling in BL conditions and emphasizing the relevance of this signalling pathway in MCs, as no other cell-type shows p-CREB induction after BL stimulus or forskolin treatment (Fig. 3C and Suppl. Fig 1A). Chromophores, which isomerize upon light absorption and drive G-protein activation, must be recycled in non-neuronal cells (retinal pigment epithelium and MCs) to sustain neuronal opsin activation ^2,15^. Indeed, when checked in the presence of 9-*cis* retinal an overall activation is observed in all of the cellular layers of the retina (Suppl. Fig. 1B). In the same line neuronal primary cultures of E8 retinas show different calcium responses when submitted to BL stimulation that are completely lost when 9-cis retinal is not added to the culture media (unpublished data). This evidence points out that in embryonic stages MCs already show the ability to specifically and autonomously respond to BL stimulation by activating CREB. Strikingly the MCs CREB activation is not uniform along the retina, somehow resembling other developmental patterns as described for retinal progenitor cells (RPCs), where they regionally gain the ability to produce specific cell fates, and lose the ability to produce others, as they bear changes in competence over time, ^51^.

Activated CREB modifies gene expression by turning on CRE-containing genes that includes immediate-early genes; metabolic regulators; survival and trophic proteins; and, particularly in the nervous system, lineage programs by controlling cell cycle progression and differentiation ^50,52–56^. In the present work we detect an increase in the percentage of cells ongoing S-phase 10 h after a single BL stimulation (Fig. 4), which can be attributed to a cell cycle lengthening rather than a S-phase arrest, given that no deleterious effect is observed in MCs cultures after the selected light protocol (suppl. fig. 2). Linking the cell cycle to cell fate decisions is a common theme in multipotent cells and has been proposed for neural progenitors as well, herein cells undergo an overall cell cycle lengthening immediately before neuronal differentiation, due to an extended G1 phase as progenitors switch from proliferative to neurogenic divisions ^57,58^. Further, we demonstrate that light-triggered cellular responses imply decreased migration and proliferation rates in MC cultures that is already evidenced by 24 h after light stimulation and sustained along time (Fig. 4). Interestingly the effect on proliferation is dependent on serum concentrations (not observed at 20% FBS media) and shows the major gap when evaluated without serum addition (0% FBS). In fact, cells illuminated and maintained in non-supplemented media (0% FBS) show no further increase in proliferation rate after 24 h, indicating light as a commanding stimuli to define their cellular fate. Most important, the effects of blue light on migration and proliferation capacity has also been detected for the opsin-expressing human MCs (MIO-M1, Fig. 4) ^22,30^, suggesting a common and conserved signalling pattern and cellular effects directed by light in MC for different species.

In retinal cultures and retinal tissue undifferentiated retinal progenitor cells preserve stem cell characteristics by expressing nestin, Sox-2, and Pax6, required for the proliferation and expansion of retinal progenitors; thus the perpetuation of these attributes is a characteristic of multipotent cells ^59–63^. In neuroespheres derived from mice cortex temporal re-localization followed by decrease in SOX2 immunolabeling is detected in cell cultures along 5 days ^64,65^, similar to the results shown in Suppl Fig 3. In our present study we evaluated the time line for nuclear SOX2 expression detecting a decrease in the percentage of SOX2-positive nuclei and decreased *PAX6* transcript by 12 and 24 h after blue light stimulation(Fig. 5). These results are in concordance with the observed effect on proliferation and directly related to cAMP signaling, given the similar effect obtained with FSK stimulation (Fig 5). The role for cAMP in differentiation processes has been described for a multiplicity of cell types including glial and neuronal types ^25,54,64,66–68^, and further it has been directed linked to CREB activation to drive astrocyte into neuronal reprogramming ^50^. An important feature of cultured embryonic Müller cells (MCs) is their ability to retain multipotency, proliferative capacity, and stem cell characteristics while preserving their retinal identity ^69^. The maintenance of these characteristics—and their specification toward MC, RPC, or neuroblast-like phenotypes—is tightly regulated by culture time, media conditions, and trophic factors ^70,71^. In line with this, evaluating our dark-adapted culture conditions across varying serum concentrations (0%, 10%, and 20% FBS) revealed that 10% FBS decreased cell migration and proliferation compared to the other conditions. Nevertheless, expression of the stem cell marker SOX2 was maintained at levels comparable to non-supplemented media (Suppl. Fig. 2). In retinal tissue *in vivo*, SOX2 is progressively downregulated during neuronal differentiation and selectively restricted to MCs, remaining absent in post-mitotic neurons ^70^. This persistent SOX2 expression indicates that while terminal neuronal differentiation entails the loss of stemness, glial populations like MCs natively preserve a multipotent molecular signature. These results support mixed populations in glial cell cultures, some maintaining multipotency while others differentiating to MCs in optimized medium conditions ^56,69,71,72^, similar to the mixed populations found within the retina in uninjured and lesioned protocols^73^.

Thus far, we have shown that a blue light-driven signaling cascade suppresses proliferation and downregulates stem cell markers in multipotent primary MCs. Therefore, we next examined which cell types are differentially represented in these cultures following illumination. We first checked for glial markers GS,RGR and GFAP and observed no-increase nor any modification on protein cellular distribution 24 h after BL stimulation (Fig. 6). We next examined the pathways across neuronal development, herein intermediate progenitors, derived from radial glial cells, are committed neurogenic progenitors, characterized for losing their proliferative capacity, decreasing PAX6 expression and specifically temporarily expressing the transcription factor Tbr2 (EOMES gene)^74,75^. This transcriptional factor is also important for retinal neurogenesis and particularly required to regulate the formation and survival of intrinsically photosensitive retinal ganglion cells, belonging to the RGC population which are the first specified cell type that emerge from a common pool of retinal progenitors ^74,76^. In our experimental conditions increased Tbr2 immunolabeling is detected 48 and 72 h after the light stimulus, with a conspicuous cytosolic vesicle-like patterning (Fig 7). This pattern is detected as well in the E8 chicken retina in cells present at the most basal portion of the developing inner nuclear layer, with no positive immunostaining in the nucleus of RGCs. This discrepancy with the expected nuclear staining for Tbr2 in the retina might indicate differences in experimental models or in developmental stages, given that RGCs are already differentiated by E8 in the chicken retina and only a small proportion (ipRGCs) retain Tbr2 expression at later stages ^76–79^. Although TBR2 is classically treated as a nuclear transcription factor, there are situations during differentiation where its localization is less strictly nuclear. Misplaced differentiating cells in the cortex show Tbr2-positive immunolabeling concentrated in the cytoplasm and in radially aligned apical processes instead of the normal nuclear distribution seen in basal progenitors of the subventricular zone. This mixed pattern suggests that when progenitors differentiate outside their usual niche, Tbr2 localization can become more diffuse across the whole cell rather than remaining confined to the nucleus, nevertheless this could be reflecting an abnormal or transitional state rather than the standard endpoint of neuronal maturation^80^. A separate study in cultured cells also reported that Tbr2-positive granules can be seen in both the nuclei and cytoplasm, whether these cells are migrating or not, respectively. This does not redefine the canonical cortical pattern, but it does support the idea that TBR2 localization can respond to cell state and environment^81^. In this direction, differentiation-linked relocalization between nucleus and cytoplasm is a common behavior that has been described for other regulatory proteins ^82^. Finally, we further confirmed that the new identity of cells in MC cultures after blue light stimulation is consistent with neuronal cell types in our culture systems, regardless of the medium conditions (Fig. 8). The increase in ß3-tubulin protein expression is a hallmark of developing and mature neurons (Suppl. Fig. 4B)^69^, and mirrors an increase in the percentage of Brn3 positive cells, an established RGCs marker ^21,83^. An interesting pattern emerges as Brn3 nuclear expression is detected in neuron-like cells, bearing small nuclei; and in the cytoplasm of MC-like cells. Brn3 ectopic expression is also cytoplasmic and not exclusively nuclear when cellular identity and developmental programs are compromised ^84^. Although nuclear trafficking mechanisms of various transcription factors seem to work differently in MC under culture conditions, still they clearly show light-directed responses. Light-guided differentiation has been reported for non-ocular cell types and directly linked to non-visual opsin activation, mainly Opn3 responding to blue and green wavelengths ^85–88^. Particularly in MC light-directed responses have been reported to drive calcium mobilization^13,14^, modify chromophore recycling^9,89^ and trigger the cell cycle re-entry and dedifferentiation of MG into a progenitor-like state ^90^. Overall, we present substantial novel evidence for an intrinsic blue light triggered signal in MC cultures that potentially drives their differentiation into retinal neurons, independently of the medium conditions, pointing out photodetection as a strong signal re-defining MC cellular fate.

## Concluding remarks

Retinal embryogenesis begins with a restricted number of radial cells, the retinal progenitor cells (RPC) and follows an hierarchical lineage model of neuronal subtype genesis where cellular subtypes are generated in a time and spatial ordered wave pattern ^51,91,92^. Retinogenesis culminates with depleted RPCs after producing five main types of neuron and the late defined MC lineage, which begin their postmitotic lives greatly resembling RPCs and possibly arising as a ‘default’ without undergoing irreversible cell fate determination^18,59,85^. Following retinal injury MC triggers differential responses across species, whereas MC to RPC conversion in regeneration-competent species resemble some developmental hallmarks, the induction of reactive gliosis by proliferation may be one of the mechanisms underlying the limited regenerative capacity of mammalian Müller glia ^63,93–95^. Light is the most powerful stimulus driving retinal function and has been recently related to non-canonical developmental configurations in cones^96^. Understanding how this extrinsic factor regulates cellular differentiation and neuronal development in the retina could pave the way for a new generation of therapeutic strategies targeting developmental and neurodegenerative disorders.

## Acknowledgments

We thank Dr. Cecilia Sanchez (CIBICI-CONCIET-UNC) and Astrid Limb for providing the MIO-M1 cell line. The authors greatly acknowledge the support staff Fabricio Navarro, Rosa Andrada, and Laura Argarañas for the care of the avian facility for the procurement of fertile eggs; Tech. Gabriela Schachner and Dr. Natalia Soledad Baez for their assistance in cell culture; the technical and imaging assistance of Dra. Cecilia Sampedro, Dr. Carlos Mas, Dra. Pilar Crespo and Dr. Gonzalo Quassollo from the Centro de Micro y Nanoscopía de Córdoba – CEMINCO – CONICET – Universidad Nacional de Córdoba, Córdoba, Argentina, (http://ceminco.conicet.unc.edu.ar); Eng. Marcelo Pino for his help in the design of irradiators for light stimulation; Daniela F. Bussolino and Vanina Torres Demichelis from the Central Facility for Media and Solutions (FACEMES). We are grateful with Dr. Lucas Sosa and Dr. Gastón Bisig, Dr. Eduardo Garbarino-Pico (CIQUIBIC-CONICET-UNC), and Dr. Jose Luis Bocco (CIBICI-CONCIET-UNC) for the kind gift of antibodies (Sox2, TBR2, Tuj-1, G3BP1 and p53) used in the present study.

---

**Supplementary Figure 1. Specific MC activation by BL stimulus implies cAMP activation in retinal explants**. BL stimulation to dark adapted retinal explants increases p-CREB expression by 30min in Vimentin positive cells (A, arrows in left image); similarly Forskolin (FSK) incubation over 30min activates p-CREB staining in Vimentin positive cells at the inner nuclear layer (INL) suggesting a sensitivy cAMP signalling in MC in E8 retinas (A, arrows in right image). When evaluated in presence of the 9-cis retinal chromophore non-specific p-CREB activation after 1h blue light (BL) stimulation is observed troughout the different retinal layers (B, arrows in right image). ONL: Outer Nucelar Layer; INL: Inner Nuclear Layer; GCL: Ganglion Cell Layer

**Supplementary Figure 2. Differential proliferative capacity in Müller glial cells cultures.** MC maintained in 10% FBS culture media display a 12h cell cycle, determined by propidium iodide flow citometry, with significant decrease in cells at the GO/G1 stage concomitant with significant increases in the percentage of S-G2/M cells by 10h (Two way ANOVA, interaction cell cycle stage*time F(2, 80)= 16.84, p<0.0001, Bonferroni posthoc test p<0.001 and p<0.01, respectively; n=10-19) (A). When tested at different serum concentration in darkness conditions (0-10-20% FBS) a significant lower rate in cell migration and proliferation is observed at 10% FBS. For wound closure % (B left graph): One way ANOVA F(2,142)= 7.21; P<0.001, Bonferroni post hoc comparison 0% vs 10% p<0.0001, n=4 independent samples. For growth rate (B right graph): Two way ANOVA RM interaction time*groups F(6, 258)=13.71; P<0.0001, Bonferroni post hoc comparison 0% vs 10% p<0.0001 at 48,72 and 96h; 10% vs 20% P<0.001 at 48 and 72h, n=4 independent samples. Noteworthy, migration and proliferative capacity does not differ in MCs cultures supplemented or not with high concentration of FBS (20%) (for wound closure %: Bonferroni post hoc comparison 0% vs 20% p>0.05. For growth rate: Bonferroni post hoc comparison 0% vs 20% p>0.05. at all of the evaluated time points).The graphs show the mean±SEM, *p<0.05 different from undifferentiated conditions. The percentage of Sox2 positive nuclei does not differ between different culture conditions (Supplemented with 0% or 10%FBS. U=110.5 p>0.05-C), nevertheless differences in Sox2 immunolabeling between the 2 conditions can be observed in the representative microphotographs as a more robust staining and wide-spread distribution in MC with 10%FBS supplemented media (D).

**Supplementary Figure 3. Blue light stimulation is innocuous to Müller glial cells in culture.** To evaluate BL-driven deleteriousa effect the transcriptional factor p53 localization was analyzed 24h after two different light intensities (microphotographs in panel C), no effect is observed at the selected light intensity (1400Lx) meanwhile positive nuclear immunostaining is observed with higher intensities (arrow at 3000Lx image). Stress granule formation was evaluated by Ras GTPase-activating protein-binding protein 1 (GRBP1) immunostaining (microphotographs in panel D). No stress granule formation is detected at the working light intensity (1400Lx), whereas protein accumlulation as cellular punctua is observed with higher intensities (arrow at 3000Lx image).

**Supplementary Figure 4. Temporal expression profile of stem cell and neuronal markers in developing neurons.** Neurospheres derived from neuronal progenitors cells from mice cortex show time-dependent modifications in SOX2 cellular localization followed by decreased expression across 6 days in optimum culture conditions (Panel A). ß3-tubulin expression by Tuj-1 clone detection, indicates positive expression from initial stages with a slight increase in immunolabeling staining as the differentiation process continues (Panel B).

**Supplementary table 1.** Summary of the antibodies used indicating the dilution for each molecular technique. ICC: immunocytochemistry; IHC: immunohistochemistry; WB: western blot; CAT#: catalogue number

