## Supplementary figures and images for "Blue light compromises cellular fate of intrinsically photosensitive Müller glial cells in the developing chicken retina"

### Supplementary Figure 1

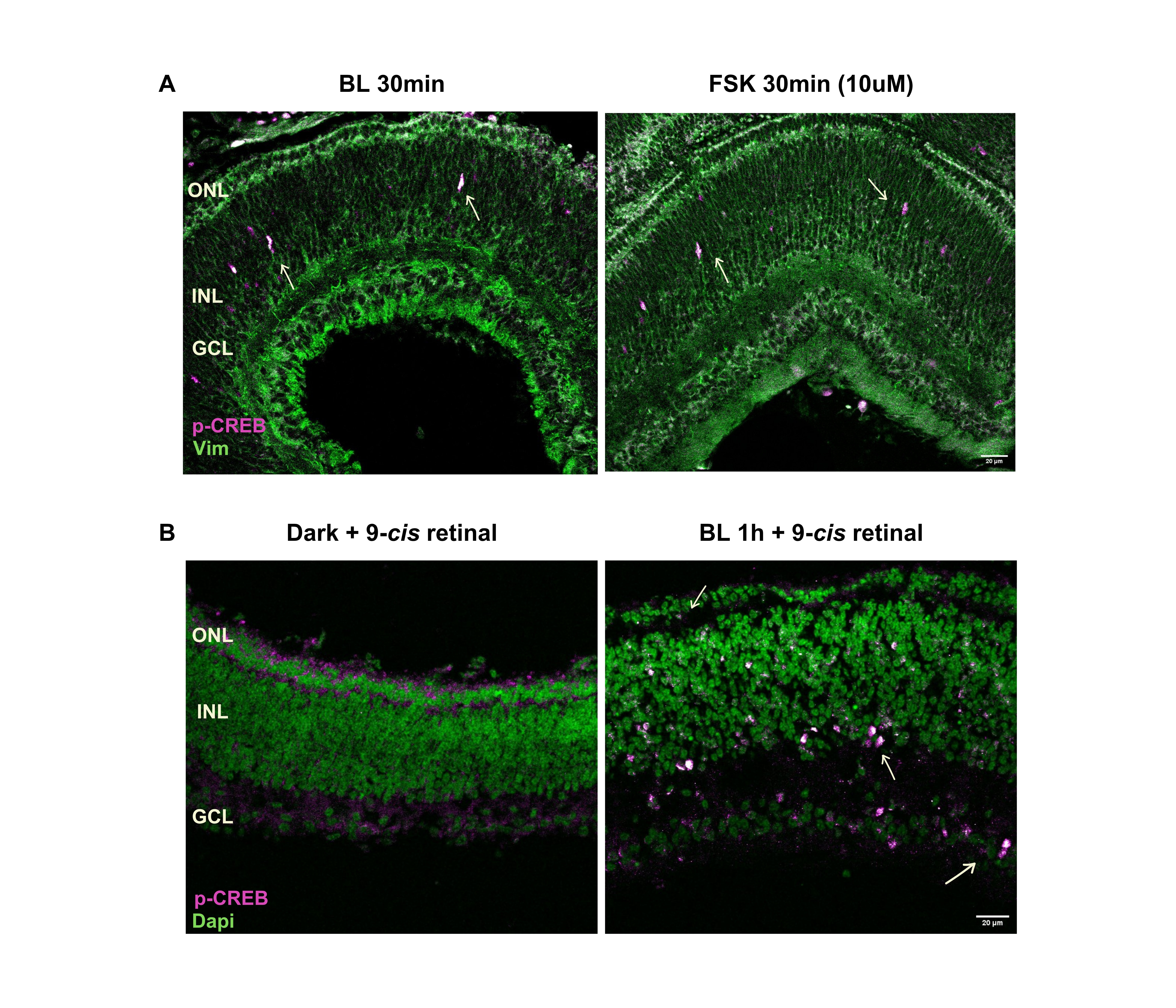

### Supplementary Figure 2

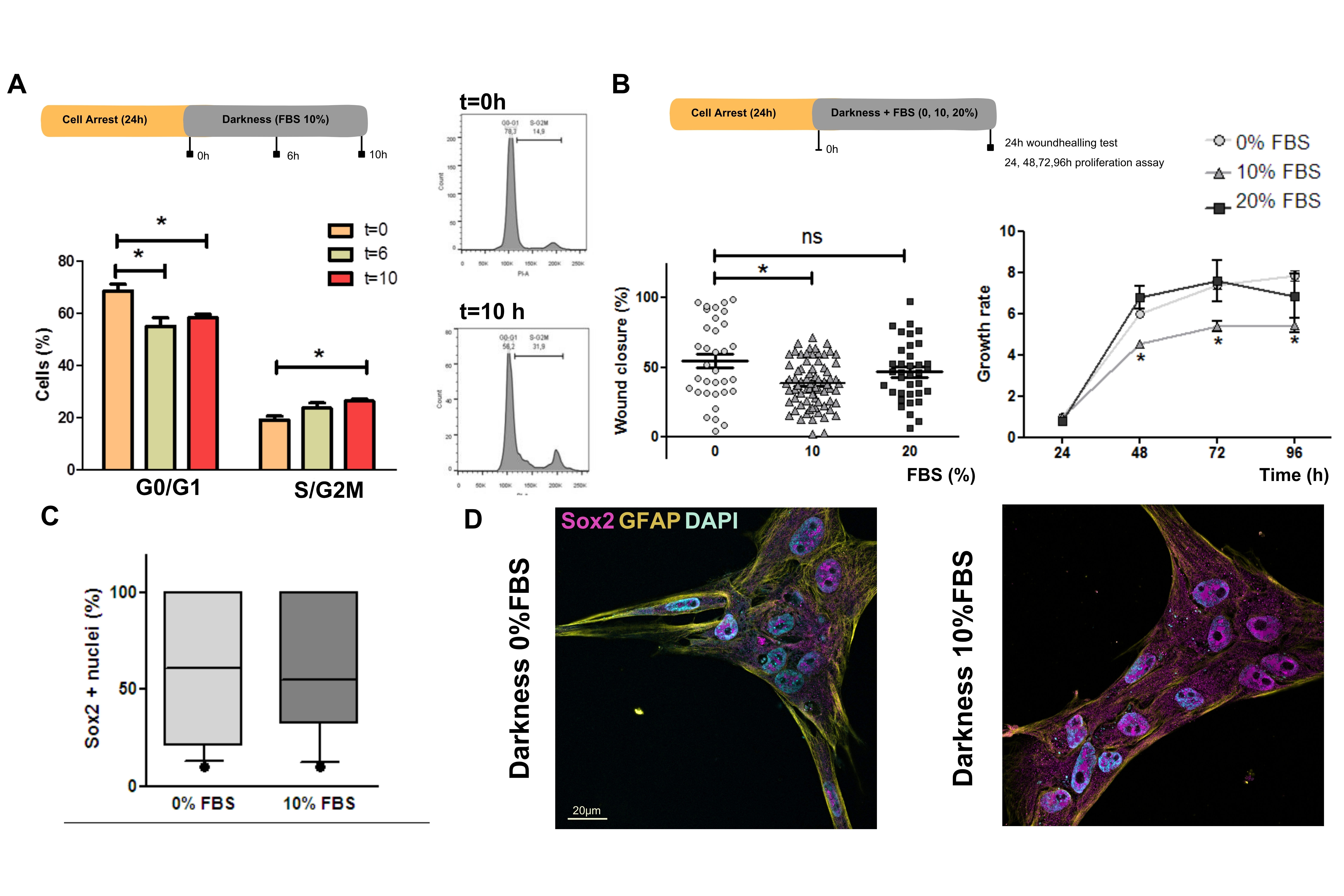

### Supplementary Figure 3

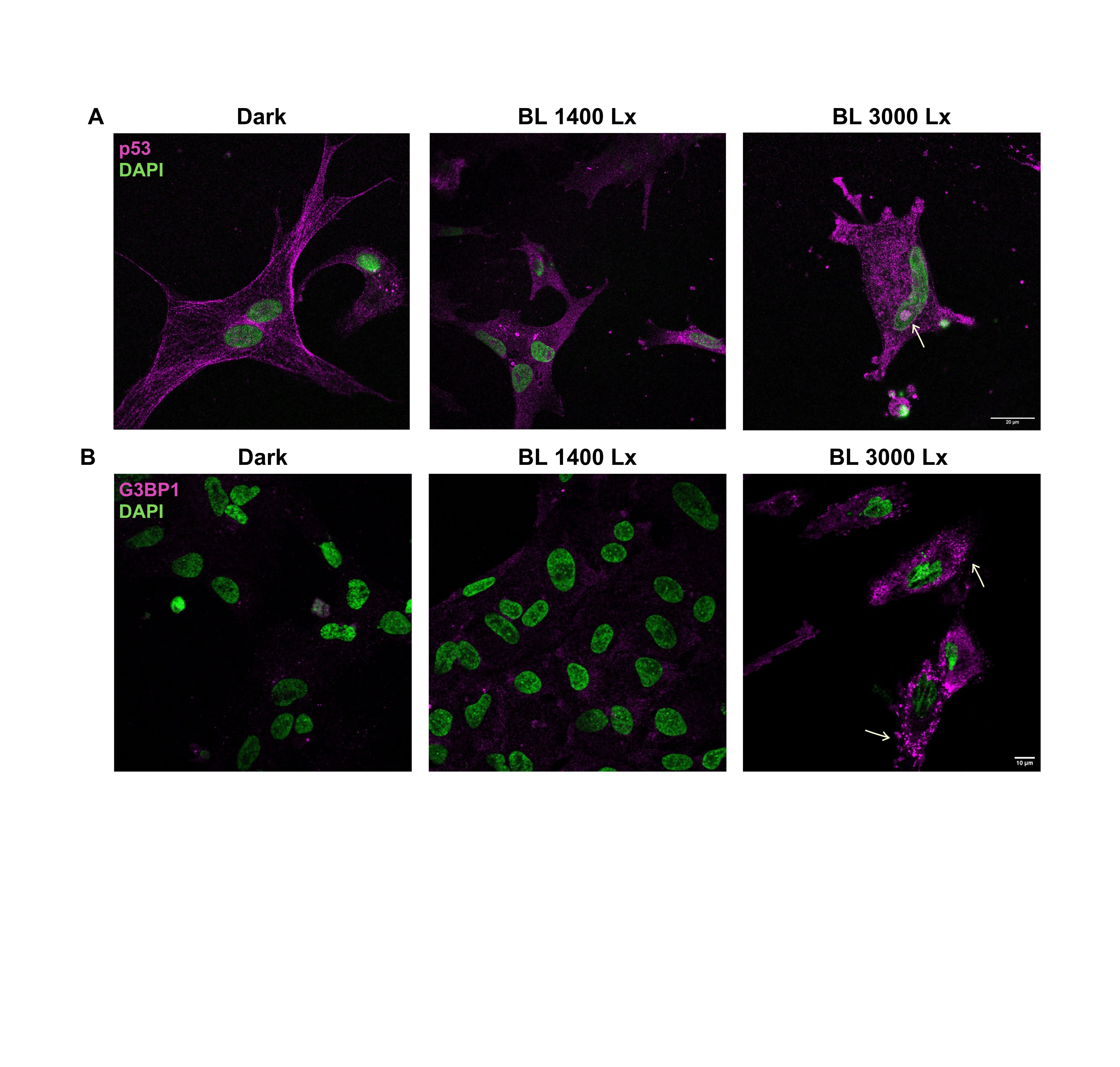

### Supplementary Figure 4

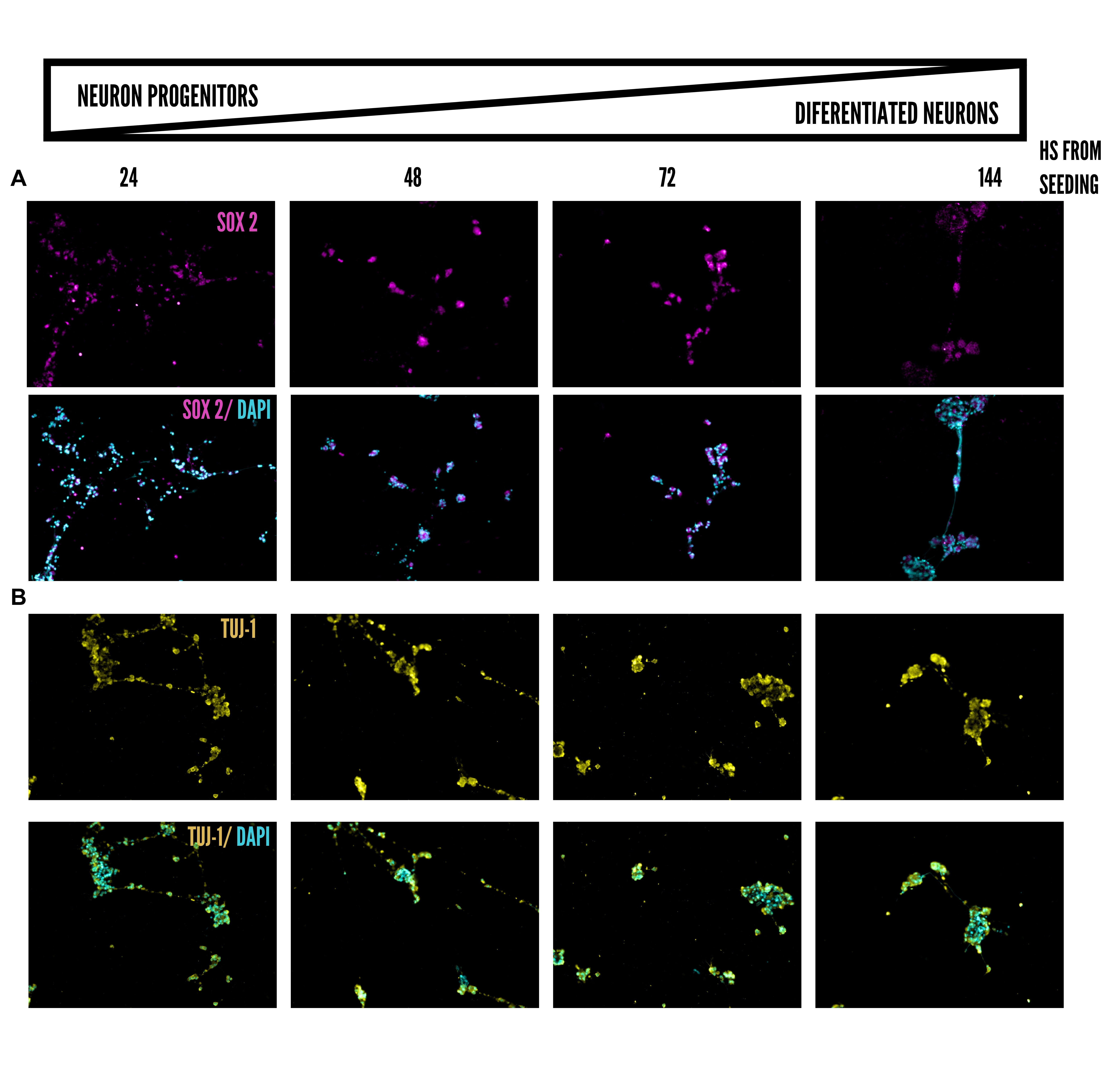
