## Supplementary Table 1 for "Blue light compromises cellular fate of intrinsically photosensitive Müller glial cells in the developing chicken retina"

| Antibody | RRID | ICC/IHC | WB |
|---|---|---|---|
| p-CREB | (Cell Signaling Technology Cat# 9198, RRID:AB_2561044) | 1:1000 | - |
| Vimentin | (Sigma-Aldrich Cat# V5255, RRID: AB_477625) | 1:1000 | - |
| SOX2 | | 1:500 | - |
| RGR | LS-A1043-50, LifeSpan BioSciences Inc. | 1:500 | 1:1000 |
| GS | (Abcam Cat# ab64613, RRID: AB_1140869) | 1:500 | 1:1000 |
| Tbr2 | | 1:500 | - |
| Brn3 | (Millipore Cat# MAB1585, RRID: AB_94166) | 1:200 | - |
| GFAP | (Sigma-Aldrich Cat# G9269, RRID:AB_477035) | 1:500 | 1:1000 |
| p53 | (Santa Cruz Biotechnology Cat# sc-126, RRID:AB_628082) | 1:100 | - |
| G3BP1 | (Bethyl Cat# A302-033A, RRID: AB_1576539) | 1:800 | - |
| tubulin | (Sigma-Aldrich Cat# T9026, RRID: AB_477593) | 1:1000 | 1:2000 |
| III-β-Tubulin | (Novus Cat# NB100-57388, RRID: AB_844201) | - | 1:500 |
| β-actin | (Sigma-Aldrich Cat# A1978, RRID: AB_476692) | - | 1:2000 |
